# Systematic evaluation of the causal relationship between DNA methylation and C-reactive protein

**DOI:** 10.1101/397836

**Authors:** Esther Walton, Gibran Hemani, Abbas Dehghan, Caroline Relton, George Davey Smith

**Affiliations:** MRC Integrative Epidemiology Unit, School of Social & Community Medicine, University of Bristol, Bristol, BS8 2BN, United Kingdom; Department of Biostatistics and Epidemiology, MRC-PHE Centre for Environment and Health, School of Public Health, Imperial College London, London, W2 1PG, United Kingdom

**Keywords:** CRP, Mendelian randomization, epigenetics, ALSPAC, inflammation

## Abstract

Elevated C-reactive protein (CRP) levels are an indicator of chronic low-grade inflammation. Epigenetic modifications, including DNA methylation, have been linked to CRP, but systematic investigations into potential underlying causal relationships have not yet been performed.

We systematically performed two-sample Mendelian randomization and colocalization analysis between CRP and DNA methylation levels, using GWAS and EWAS summary statistics as well as individual level data available through the ARIES subset of the Avon Longitudinal Study of Parents and Children (ALSPAC; 1,616 participants).

We found no convincing examples for a causal association from CRP to DNA methylation. Testing for the reverse (a putative causal effect of DNA methylation on CRP), we found three CpG sites that had shared genetic effects with CRP levels after correcting for multiple testing (cg26470501 (offspring: beta=0.07 [0.03, 0.11]; mothers: beta=0.08 [0.04, 0.13]), cg27023597 (offspring: beta=0.18 [0.10, 0.25]; mothers: beta=0.20 [0.12, 0.28]) and cg12054453 (offspring: beta=0.09 [0.05, 0.13])) influenced CRP levels. For all three CpG sites, linked to the genes *TMEM49, BCL3* and *MIR21*, increased methylation related to an increase in CRP levels. Two CpGs (cg27023597 and cg12054453) were influenced by SNPs in genomic regions that had not previously been implicated in CRP GWASs, implicating them as novel genetic associations.

Overall, our findings suggest that CRP associations with DNA methylation are more likely to be driven by either confounding or causal influences of DNA methylation on CRP levels, rather than the reverse.

## Introduction

C-reactive protein (CRP) is a marker of chronic low-grade inflammation and high CRP levels are linked to increased risk of mortality (1) and a range of disorders (2), including coronary heart disease and stroke (3). The largest genome-wide association study (GWAS) on CRP levels to date identified 18 independent genetic loci (4). However, several other molecular mechanisms such as epigenetic modifications, which are downstream of genetic variation, might also be involved in CRP functioning.

Epigenetic modifications, including DNA methylation, can affect gene expression (5) without changing the DNA sequence. To unravel the role of DNA methylation in chronic inflammation, Ligthart et al. (6) performed an epigenome-wide association study (EWAS) on serum CRP levels and identified 58 DNA methylation markers in a large panel of individuals of European descent (n=8,863) and replicated in a panel of African Americans (n=4,111). However, currently it is not well understood whether DNA methylation is a cause or consequence of changes in CRP levels.

In Mendelian randomization (MR) genetic variants are used as instrumental variables to assess the causality of an association (7–9). Using genetic variants as instruments can reduce biases, which arise due to confounding or reverse causation. For an instrument to be valid, it must be i) reliably associated with the exposure; ii) associated with the outcome only through the exposure and iii) independent of confounders that influence the exposure and the outcome. To address the current question, we made use of genetic variants that act independently as proxies for CRP levels and for DNA methylation markers.

With respect to DNA methylation data, an MR framework has been proposed (Relton & Davey Smith 2012) and has, for instance, been applied to investigate the causal relationship between *HIF3A*-linked DNA methylation and body mass index. In that study, the authors failed to identify a causal effect of DNA methylation on body mass index (10). With relevance to CRP and epigenome-wide data, a recent epigenome-wide association study identified CpG sites that putatively influenced cardiovascular disease risk traits, including CRP (11). The authors reported that some cardiovascular traits are influenced by altered DNA methylation, but evidence specific to CRP was weak.

In this study, we applied MR to infer causality and the direction of effect between CRP and epigenome-wide DNA methylation markers.

## Results

### Causal effect of CRP on epigenome-wide DNA methylation

To investigate whether CRP is causally linked to DNA methylation at an epigenome-wide level, we used 18 SNPs, which have been shown to associate with CRP levels (4), as genetic instruments. Genotype-exposure estimates were taken from the same study. To derive genotype-outcome estimates, the effect of these 18 SNPs on epigenome-wide DNA methylation was tested in a sample of 1,616 participants from the ARIES study (see “MR sample” in Table 1).

**Table 1.**
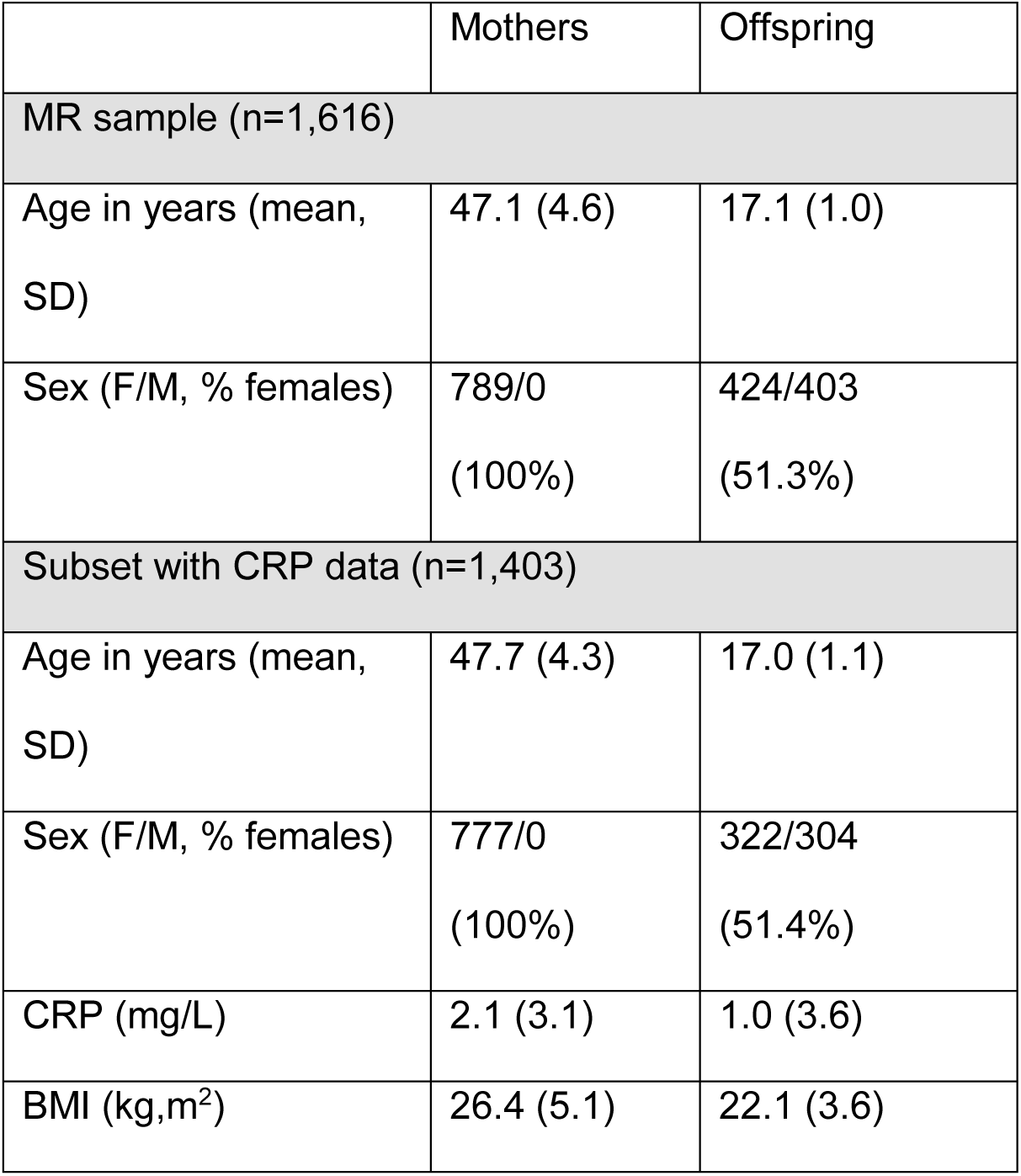
ARIES study sample characteristics.

|  | Mothers | Offspring |
| --- | --- | --- |
| MR sample (n=1,616) |  |  |
| Age in years (mean, SD) | 47.1 (4.6) | 17.1 (1.0) |
| Sex (F/M, % females) | 789/0<br>(100%) | 424/403<br>(51.3%) |
| Subset with CRP data (n=1,403) |  |  |
| Age in years (mean, SD) | 47.7 (4.3) | 17.0 (1.1) |
| Sex (F/M, % females) | 777/0<br>(100%) | 322/304<br>(51.4%) |
| CRP (mg/L) | 2.1 (3.1) | 1.0 (3.6) |
| BMI (kg,m<sup>2</sup>) | 26.4 (5.1) | 22.1 (3.6) |

Using a range of MR methods, including IVW, WM and MR-Egger, to combine the effect from individual SNPS, we found no evidence for an association with single site DNA methylation at an epigenome-wide threshold (Figure 2 and Supplementary Material (SM) Table S1). A lambda of 0.99 and the inspection of the QQ plot (SM S1A) provided little evidence for population sub-structures or genotyping errors. We also failed to identify differentially methylated regions (i.e., a region of the genome in which several neighbouring CpG sites show differential methylation, SM Figure S1B, and SM Figure S2 for CpG-specific regional plots and gene track annotation). Furthermore, we found no evidence for causal associations restricting ourselves to the 58 CpG sites that were linked to CRP in a previous EWAS (6). In detail, there was no correlation between effect sizes of the EWAS and MR analyses and no CpG site passed a threshold correction for 58 tests (SM Table S2 and SM Figure S3 for CpG-specific regional plots).

**Figure 1.**
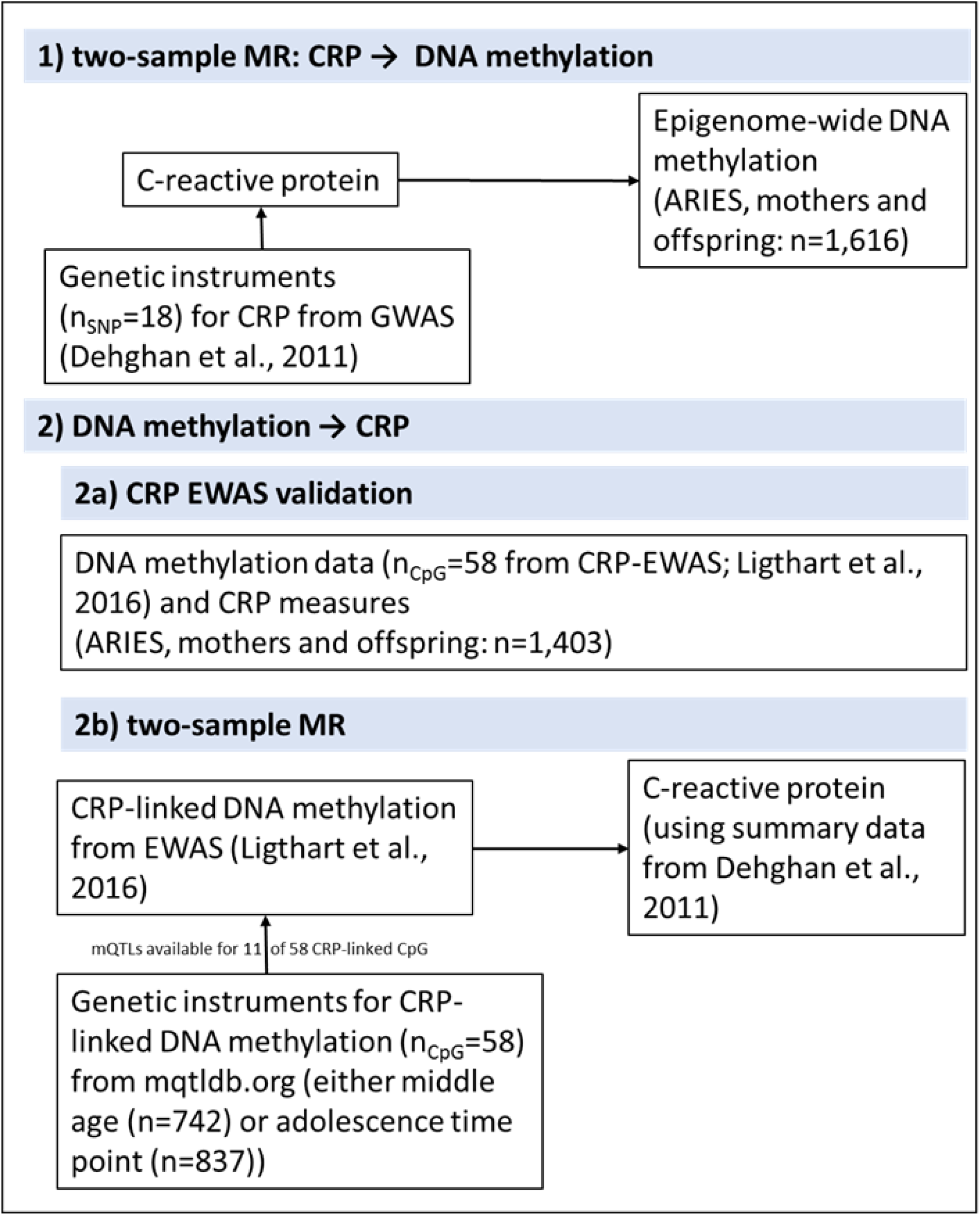
Method overview.

**Figure 2.**
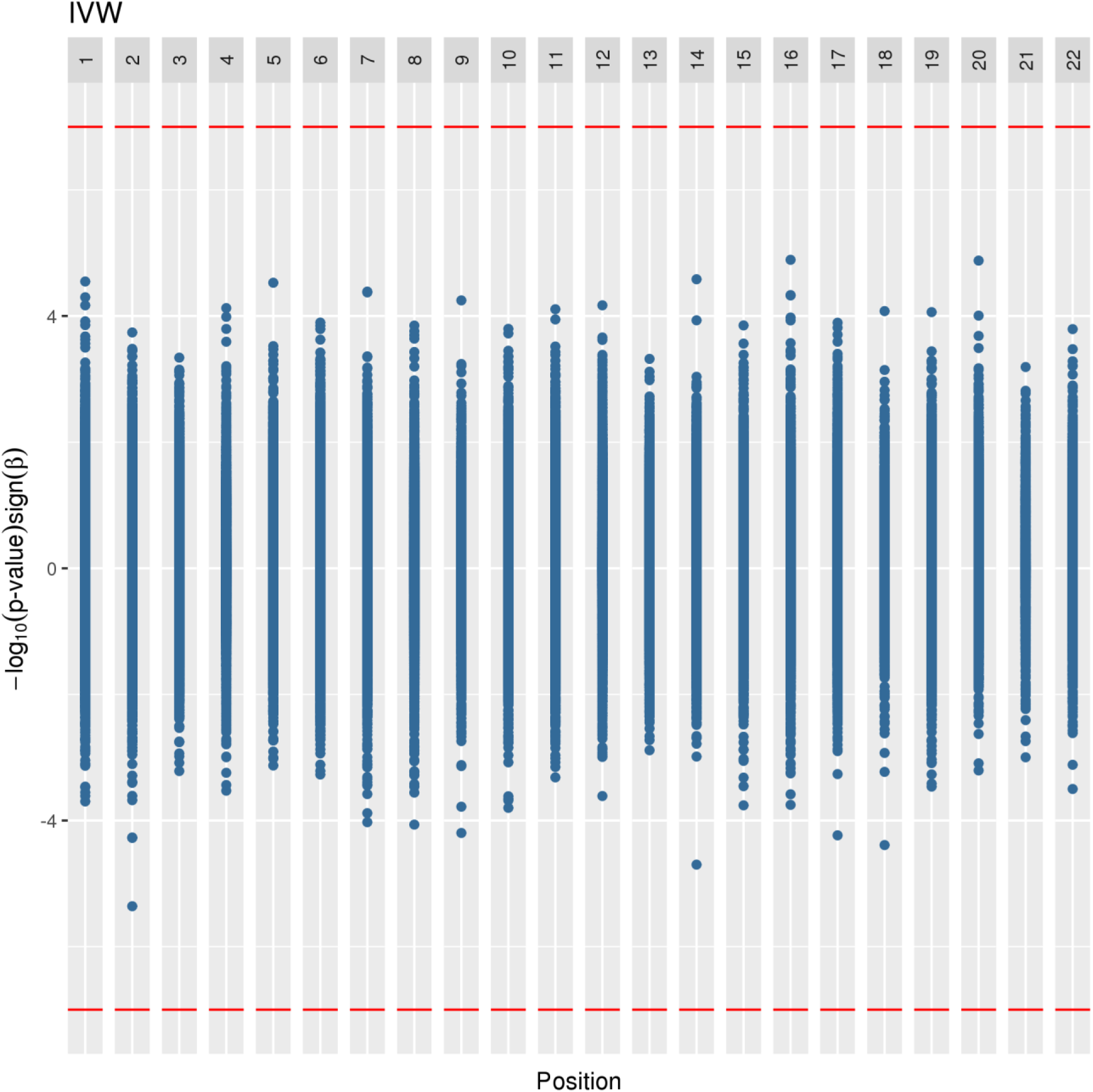
Causal relationship of CRP on epigenome-wide DNA methylation. CpG position in the genome (x-axis) is plotted against −log(p-values) multiplied by the sign of the MR estimate (y-axis). IVW, inverse-variance weighted method.

### Replication of CRP-linked DNA methylation effects on CRP

As a validation analysis, we first sought to confirm previous CRP EWAS findings in our dataset (see “Subset with CRP data” in Table 1). We identified a strong correlation between our and previously reported effect sizes of the 58 CpGs sites (rho = 0.83, *p* = 9.49^∗^10^−16^, 95% CI [0.73, 0.90]; SM Figure S4). For 55 of 58 CpGs, the sign of effect was consistent between the Ligthart et al. (6) results and our own (SM Table S2).

### Causal relationship of CRP-linked DNA methylation on CRP

As a next step, we tested for a causal effect of DNA methylation on CRP, separately in the mothers and offspring, as mQTLs were identified specific to each group and time point. mQTLs could be identified (via mqtldb.org) for 13 of the 58 CpG sites that were originally identified to be linked to CRP. mQTLs for 11 of these 13 CpGs, linked to eight genes, were available in the summary data from Dehghan et al. (4) and subsequently analysed in an MR framework (see SM Figure S5 for a flowchart and further details). In the mothers, one CpG (cg08548559) was tagged by two independent mQTLs, while in both mothers and offspring one mQTL (rs12677618) tagged two different CpG sites. For cg12054453, we could identify an mQTL in the offspring only. MR analysis indicated that three CRP-linked CpG sites (cg26470501, cg27023597 and cg12054453, for which an mQTL was only detectable in the offspring) were putatively causal for CRP levels. Two of these three CpG sites are located on chromosome 17 within 2,545 base pairs from each other and link to the genes *Transmembrane Protein 49* (*TMEM49*) and *MicroRNA 21* (*MIR21*). The third CpG lies on chromosome 19 and is linked to the gene *B-Cell Lymphoma 3 Protein* (*BCL3*). While this CpG is located 170 kbp away from one of the 18 CRP GWAS loci (rs4420638), the two CpGs on chromosome 17 are not located near any of the previously reported GWAS findings suggesting a potentially novel genomic region to be linked to CRP. For all three CpG sites, increased methylation related to an increase in CRP levels (Figure 3 and SM Table S3). The sign of the putative causal effects for these three CpG sites was opposite to the observed associations reported in the current study and in Ligthart et al. (6).

**Figure 3.**
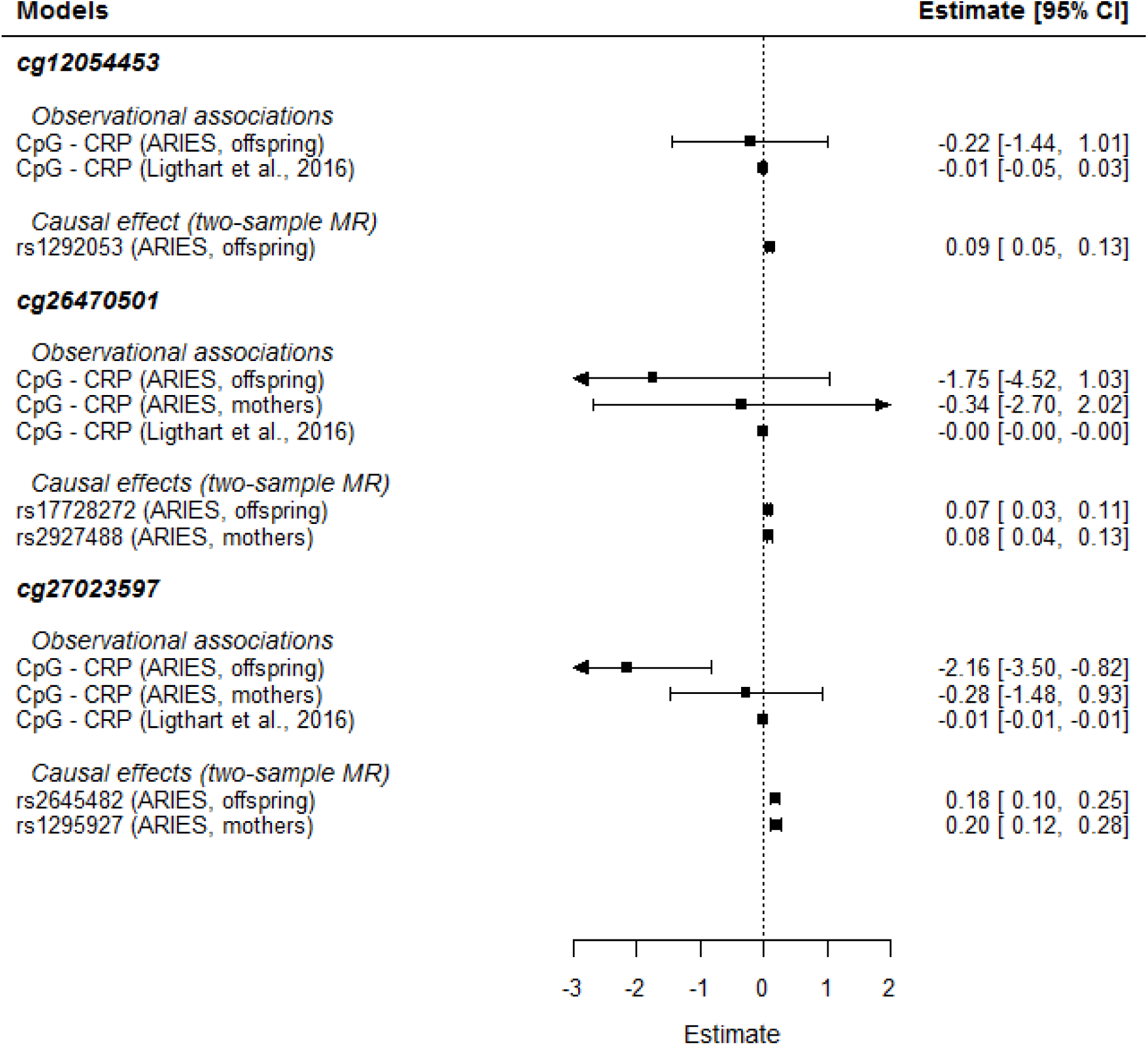
Observational and causal relationship of CRP-linked DNA methylation on CRP based on a two-sample MR. An mQTL for cg12054453 was only available in the offspring.

### Additional analyses

The three putatively causal CpG sites were each instrumented by only a single genetic variant. It is hence possible that such an mQTL is merely in linkage disequilibrium with another variant that affects CRP levels. To address this, we carried out a colocalization test for each CpG and all SNPs within a 1MB window of the instrumental mQTL. In each case, results provided strong evidence for one shared causal variant, given posterior probabilities for this scenario between 77.4 – 98.2% (SM Table S7). Of note, these analyses were based on a small number of SNPs (range: 2-57) and did not allow us to distinguish between mediation (SNP to methylation to CRP) and horizontal pleiotropy (SNP affecting methylation and CRP via independent biological pathways).

As CRP is predominantly expressed in the liver, but DNA methylation was obtained from blood cells. We were interested to see whether the three genes linked to the putatively causal CpG sites were expressed more strongly in liver tissue compared to the remaining five genes with no causal CpG sites. We therefore investigated gene expression profiles of all eight genes that linked to the investigated 11 CpGs, using public data available via gtexportal.org. Gene expression of TMEM49 (median transcript per million (TPM) = 29.54) and BCL3 (median TPM = 100.40) was highly elevated in the liver compared to the other genes (range of median TPM: 0 – 4.24; SM Figure S6), suggesting a relevance for these genes in liver tissue.

As a last step, we investigated whether the five SNPs, which were used as instruments for the three putatively causal CpG sites, were also linked to BMI, as i) BMI and CRP are closely linked (12) and ii) BMI has been shown to influence DNA methylation (13). An association of the instrumental SNPs with BMI could indicate a potential violation of MR assumptions; however, a look-up of these SNPs in a recent GWAS on BMI (14) provided no clear evidence associations (SM Table S8).

## Discussion

Applying MR to infer causality and the direction of effect between CRP and epigenome-wide DNA methylation, we found little evidence that CRP causally impacts DNA methylation levels. Instead, for a selection of CpG sites, DNA methylation appeared to influence CRP levels. For these sites, which were linked to the genes *TMEM49, BCL3* and *MIR21*, the putative causal effect on CRP was opposite in sign to the observed associations, suggestive of potential negative feedback mechanisms.

Several studies have reported associations between DNA methylation — either in candidate genes such as *AIM2* or epigenome-wide — and CRP levels (6, 11, 15). Most notably, the largest epigenome-wide meta-analysis so far identified and replicated 58 CpG sites. Interestingly, while 51 of these 58 sites were also linked to at least one cardiometabolic trait, only 9 sites were also associated with whole blood gene expression levels in cis. While the authors were careful to not draw conclusions about the causality of effects, these results suggest that the route from DNA methylation to gene expression and disease might be more complex in some contexts. Miller et al. (15) followed up on these findings and focussed in particular on *AIM2* DNA methylation in relation to CRP in a sample of US military veterans with and without post-traumatic stress disorder (PTSD). They found that *AIM2* DNA methylation was lower in PTSD cases and also negatively correlated with CRP levels, which itself was elevated in patients. Specifically, the authors also reported an indirect effect of PTSD on CRP via *AIM2* DNA methylation, indicating that in the case of *AIM2*, DNA methylation might be upstream of CRP. We were not able to investigate the causal effects of *AIM2* DNA methylation in our study, as we were unable to identify an mQTL for the underlying CpG. However, consistent with both previous studies (6, 15), AIM2_cg10636246_ DNA methylation was negatively associated with CRP in our study, providing an overall agreement in observational effects between studies. Additionally, in line with Miller et al. (15), we found that for a selection of CpG sites, DNA methylation appeared to influence CRP levels — rather than the reverse. Similarly, Richardson et al. (11) reported that some genetic influences on cardiovascular traits were putatively mediated through changes in DNA methylation, which itself displayed downstream effects on gene expression. However, in that study the authors employed a one-sample MR approach that can be less powerful than a two-sample MR analysis and indeed, the evidence specifically to CRP was weak, not passing the genome-wide threshold applied in the current study. Considering that DNA methylation appeared to influence CRP levels in only a selection of CpG sites and that not all of these CpG sites were among the most strongly associated sites in the original EWAS indicates that the suggestive pathway from DNA methylation to CRP might be CpG-specific and not a general rule.

Our findings add to several studies that focussed on the interplay between DNA methylation, gene expression and protein levels. Ng et al. (16) tested for associations between epigenetic markers (including DNA methylation and histone modifications) and gene expression and found that 9% of association sets followed an epigenetic mediation model, which was defined by a propagation path from genetic variants to gene expression via epigenetic modification. Only 3% of associations showed evidence for the reverse path, where gene expression impacts the epigenome.

Contrary to this report, Ahsan et al. (17) investigated the interplay between DNA methylation and 121 protein biomarkers for inflammation, cancer and cardiovascular disease (although not CRP). They found that for 22% (i.e., n=27) of the studied proteins, biomarker levels appeared to influence DNA methylation, rather than the reserve. The authors hence argued that genetic variants, environmental exposures or secondary effects of disease impact DNA methylation, which itself might not play a direct role in the regulation of these biomarkers. Taken together with the current results, findings suggest that DNA methylation in certain genes may lie upstream of gene expression and protein levels of some inflammation-related biomarkers such as CRP, but that this finding cannot be generalized overall.

For three CpGs, we identified putative causal effects on CRP levels. Two CpGs were within 2,500 base pairs from each other, located in the proximity of the genes *TMEM49* and *MIR21*. TMEM49 is a transmembrane protein, which is expressed in most tissues, and plays a role in autophagy, an intracellular degradation process. *TMEM49* has been linked to pancreatitis, apoptosis and cell-cell adhesion. Aside its association with CRP (6), TMEM49 DNA methylation associated with two inflammatory markers (soluble tumor necrosis factor receptor 2 and — to a lesser degree — interleukin-6) prior to and 6 months after radiation therapy in a small sample of patients with breast cancer (18). Similarly, expression of miR-21, located in the intron of *TMEM49* (19), has been observed to be responsive to inflammation (20–22).

*BCL3* is a proto-oncogene candidate. It has been associated with anti-inflammatory response in diseased pancreatic rodent and human tissues (23) and BCL3 suppression increases the release of the pro-inflammatory cytokines IL-8 and IL-17 in T cell lymphoma (24). However, the link between BCL3 DNA methylation and CRP levels, which was reported in Ligthart et al. (6), could not be replicated in a recent validation study using four cohorts with sample sizes between 286 and 1,741, although in all four cohorts the sign of the association was negative (25), consistent with Ligthart et al. (6) and findings reported in the current study.

In summary, our findings further support the role of DNA methylation linked to the genes *TMEM49, MIR21* and *BCL3* in inflammatory processes. Their function in inflammation is most likely through an upregulating effect on serum CRP.

Of note, the sign of the putative causal effects identified in the current study were opposite in sign to the observational associations reported both in our study as well as in previous studies (6, 25). Discrepancies between observational and causal associations have been reported repeatedly in the literature. Much attention was given to the findings that vitamin ß-carotene deficiency apparently does not increase the risk of smoking-related cancers (26) or that there does not appear to be a causal link between vitamin C or E and coronary heart disease (27, 28), despite promising observational associations in all three cases. Measurement error and residual confounding are just two reasons that can explain this discrepancy and are discussed at length elsewhere (7). It is however noteworthy that in the case of DNA methylation (linked to *TMEM49, MIR21* and *BCL3*) and CRP, observational findings between studies were in agreement despite cross-study differences in demographics (the Ligthart et al. cohorts had a larger male:female ratio and were on average older than our cohort), in CRP levels (which were clinically elevated in Ligthart et al., but not in our study) or clinical sample characteristics (on average, rates of diabetes, coronary heart disease or hypertension were higher in Ligthart et al. compared to our cohort). Cell count differences were controlled for in both analyses, which were predominately based on data from whole blood, but it could be argued that altered blood cell type composition is part of the innate immune system’s response to inflammation and hence on the causal pathway. In fact, natural killer and CD8T cell counts have been observed to be inversely associated with CRP levels (15) and collider bias — arising when cell type composition is a function of both DNA methylation and CRP — could explain the discrepancy in findings (SM Figure S7). However, for one it seems unlikely that DNA methylation impacts cell type composition. Rather, DNA methylation is an indicator of cell type composition. Second, Ligthart et al. (6) carried out sensitivity analyses in CD4+ cells only and confirmed the association with serum CRP for 18 of the original 58 CpGs. While none of the three CpGs linked to *TMEM49, MIR21* or *BCL3* associated with CRP in this subsample, the sign of effect observed in CD4+ cell was consistent with the original findings, which were based on whole-blood tissue. It is possible that a negative feedback mechanism could account for the discrepancy in effect signs. In detail, it is conceivable that DNA methylation in the reported genes lead to an upregulation of CRP levels (i.e., the causal effect), which also results in a cascade of inflammatory-related processes (such as altered natural killer cell or CD8T cell counts) that might affect measurements of DNA methylation in whole blood (i.e., the observed effect). However, further studies are necessary to draw more definite conclusions.

The findings of this study should be considered in light of the following limitations. First, although we employed a two-sample MR setting to benefit from findings based on large, well-powered studies, mQTLs were identified in a sample of 837 adolescents and 742 mothers in middle age. It is hence possible that the mQTLs used in our study represent less optimal instruments for DNA methylation. In fact, only four of the 11 mQTLs used in our study (and none of the three CpGs linked to *TMEM49, MIR21* or *BCL3*) overlapped with those identified in Ligthart et al. (6) using data from the Framingham Heart and the Rotterdam study. It remains to be seen how well our findings replicate using mQTLs that are based on larger samples.

Second, violations of MR assumptions such as horizontal pleiotropy (i.e., variants influencing CRP through other pathways than the CpG) can limit the conclusions drawn in this study. In the case of DNA methylation, we had only one instrument available, which restricted our ability to evaluate bias due to horizontal pleiotropy. However, to reduce the risk of horizontal pleiotropy we used mainly cis-regulatory SNPs (only two mQTLs were more than 1 Mb away from their CpG). This greatly reduces the likelihood of pleiotropic effects. We also conducted colocalization analysis for putative MR associations, and while not sufficient, demonstrating shared genetic variants between a putative exposure and outcome is necessary for a causal association.

Third, we did not investigate other markers of inflammation. For instance, biomarkers upstream of CRP include IL-6 (29), which stimulates the production of CRP. A small epigenome-wide study (30) found that 25 of the 58 sites reported by Ligthart et al. (6) also nominally associated with inflammatory cytokines such as TNF, IL-6, IL-8 and IL-10. Reassuringly, the majority of these CpGs exhibited similar observed relationships (ie. positive or negative correlation) as previously reported for CRP, but an investigation of causal associations is still lacking.

In summary, we found no evidence that CRP causally impacts DNA methylation levels. Instead, for a selection of CpG sites, DNA methylation appeared to influence CRP levels in a direction that was opposite to the observed association. Our findings further support previous findings that see DNA methylation as one factor on the causal pathway to inflammation.

## Materials and Methods

### Overview

We carried out a two-sample MR analysis (31, 32) using summary statistics of GWAS and EWAS analyses on CRP as well as CRP, genome- and epigenome-wide data available through the ARIES subset of the Avon Longitudinal Study of Parents and Children (ALSPAC; (33); Figure 1). Specifically, to investigate whether CRP causes changes in DNA methylation on an epigenome-wide level (panel 1 in Figure 1), we used genetic variants (n=18) that have been reported to be robustly associated with CRP levels (4) as proxies for CRP levels and explored their effect on epigenome-wide DNA methylation using the ARIES resource (see below for further details). To examine the reverse association (DNA methylation causally influencing CRP levels; panel 2 in Figure 1), we studied CpG sites (n=58) that have been linked to CRP levels in a recent EWAS (6). We first investigated how well these previous findings could be replicated in our cohort by performing an association analysis on these CpG sites and CRP measures (panel 2a in Figure 1). Then — to investigate potentially causal effects — we identified underlying methylation quantitative trait loci (mQTLs) for these CpG sites via mqtldb.org (see below for further details) and combined this with CRP GWAS summary data (4) to inspect causal pathways within a two-sample MR framework (panel 2b in Figure 1).

### Genetic instruments for CRP levels

Summary statistics from the to-date largest GWAS of CRP levels were based on an analysis of 66,185 participants from 15 population-based cohort studies with a replication in 16,540 participants from ten independent studies (4). All participants were of European ancestry. Genome-wide scans on natural log-transformed CRP levels (mg/L) were run controlling for age, sex as well as site and family structure, if necessary. Results were subsequently corrected for genomic inflation. Eighteen SNPs passed genome-wide threshold and were used as genetic instruments for CRP in the current analysis.

### Association of 18 CRP index genetic variants with epigenome-wide DNA methylation Sample

We used CRP, genetic and epigenetic data from participants, which were drawn from the Accessible Resource for Integrated Epigenomics Studies (ARIES, http://www.ariesepigenomics.org.uk; (33)), a subset of 1018 mother-offspring pairs nested within the Avon Longitudinal Study of Parents and Children (ALSPAC; (34, 35)). ALSPAC is an ongoing epidemiological study of children born from 14,541 pregnant women residing in Avon, UK, with an expected delivery date between April 1991 and December 1992. Informed consent was obtained from all ALSPAC participants and ethical approval was obtained from the ALSPAC Law and Ethics Committee as well as Local Research Committees. Please note that the study website contains details of all the data that is available through a fully searchable data dictionary and a variable search tool: http://www.bris.ac.uk/alspac/researchers/our-data/.

### Measures

Peripheral blood samples were collected and analysed from mothers and children at the time, when the children were adolescents (mean age_offspring_ = 17.1, mean age_mothers_ = 47.7). DNA methylation data was measured using the Illumina Infinium HumanMethylation450K BeadChip. Samples failing quality control (average probe P value ≥0.05, those with sex or genotype mismatches) were excluded from further analysis and scheduled for repeat assay, and probes that contained <95% of signals detectable above background signal (detection P value <0.05) were excluded from analysis (n=1,113) leaving 481,742 CpGs to be analysed. Methylation data were pre-processed using R software, with background correction and functional normalization performed using the meffil pipeline (36).

For CRP measurements, blood samples were collected from participants who were instructed to fast overnight if their assessment was in the morning, or at least for 6 h if the assessment was in the afternoon. Blood samples were immediately spun, frozen and stored at −80 degrees Celsius, and analysed within 3-9 months of blood sampling with no freeze-thaw cycles in between. High sensitivity CRP was measured by automated particle-enhanced immunoturbidimetric assay (Roche UK, Welwyn Garden City, UK).

Genetic data was based on the Illumina HumanHap550 quad genome-wide SNP genotyping platform (Illumina Inc, San Diego, USA). Imputation was performed using a joint reference panel using variants discovered through whole genome sequencing in the UK10K project along with known variants taken from the 1000 genomes reference panel. Novel functionality was developed in IMPUTE2 (37) to use each reference panel to impute missing variants in their counterparts before ultimately combining them together. All variants were filtered to have Hardy-Weinberg equilibrium P > 5×10-7, a minor allele frequency ≥ 0.01 and an imputation quality score ≥ 0.8.

To derive effect estimates and standard errors of 18 CRP-linked SNPs (4) on epigenome-wide DNA methylation, we used data in all mothers and children with genome-wide and epigenome-wide data available (n=1,616). Using linear mixed-effects models, CpG level methylation was regressed against 18 SNPs, controlling for sex, age, cell type as well as batch and family (both modelled as random effects). To account for ancestry and in line with Dehghan et al. (4), we additionally included 10 genetic principal components. Effect estimates and standard errors were carried forward to be used in a Mendelian randomization analysis.

### Replication of CRP EWAS findings

Fifty-eight CRP-linked CpGs at a genome-wide threshold were identified in a meta-analysis (6) of a study in a large European population (n = 8,863) and a trans-ethnic replication in African Americans (n = 4,111).

To verify how well these recent CRP EWAS findings could be replicated in our cohort, we performed an association analysis on these CpG sites in the mothers and children for which CRP measures were available (n=1,403). We followed the analysis steps as laid out in Ligthart et al. (6) as closely as possible. In detail, linear mixed-effects models were applied (via the R package lme4), in which CpG level methylation (untransformed beta-values) was regressed against natural log-transformed CRP levels with adjustment for sex, age, cell type, BMI as well as batch and family (both modelled as random effects).

### mQTLs for CRP-linked DNA methylation

mQTLs for these 58 CpGs were identified using the mQTLdb database (http://www.mqtldb.org/; date last accessed: June 22, 2017), an open-access repository of methylation quantitative trait loci (mQTLs) associated with methylation levels of blood samples across the genome. mQTLs were carried forward that associated with methylation levels in ALSPAC mothers or offspring at any of the 58 CRP-linked CpG sites within a 1Mb based on exact linear regression in PLINK. We could identify mQTLs for 13 of the original 58 CpGs. mQTLs that were linked to the same CpG were subsequently pruned using the LD-clumping function in MR-Base (38). For two CpGs, none of the identified mQTLS were present in the CRP GWAS summary data, leaving 11 CpGs in the final analysis.

### Mendelian randomisation analysis

All summary statistics were harmonized using the TwoSampleMR R package, which is part of the MR-Base online platform (38). Two-sample MR analyses were then undertaken using the TwoSampleMR and MendelianRandomization R packages (39). Linear regression analysis of SNP-CRP and SNP-CpG association were carried out using an inverse-variance weighted (IVW), weighted median (WM) and MR-Egger methods. To identify differentially methylated regions (DMRs), we used the Python module comb-p (40), which is designed to identify peaks in a vector of spatially correlated P values by grouping correlated CpGs in a 500-bp sliding window with a stepsize of 50.

To test for causal effects of CRP-linked DNA methylation on CRP, we applied the Wald ratio method (41), as for most CpGs only one mQTL was available for analysis after LD pruning and data harmonization. Colocalization tests were carried out for fine mapping purposes using the coloc.abf function in the coloc R package (42). This allowed us to examine whether mQTLs were in linkage disequilibrium (LD) with other SNPs that might impact CRP levels.

## Funding

This work was supported by the UK Medical Research Council and Wellcome [Grant number 102215/2/13/2] and the University of Bristol provide core support for ALSPAC. This publication is the work of the authors and E Walton will serve as guarantors for the contents of this paper. A comprehensive list of grants funding is available on the ALSPAC website (http://www.bristol.ac.uk/alspac/external/documents/grant-acknowledgements.pdf). This research was specifically funded British Heart Foundation [grant number SP/07/008/24066] and National Institutes of Health [grant number R01 DK077659]. The Accessible Resource for Integrated Epigenomics Studies (ARIES) was funded by the UK Biotechnology and Biological Sciences Research Council [grant number BB/I025751/1 and BB/I025263/1]. Supplementary funding to generate DNA methylation data which is included in ARIES has been obtained from the MRC, ESRC, NIH and other sources. ARIES is maintained under the auspices of the MRC Integrative Epidemiology Unit at the University of Bristol [grant numbers MC_UU_12013/2 and MC_UU_12013/8]. GWAS data was generated by Sample Logistics and Genotyping Facilities at Wellcome Sanger Institute and LabCorp (Laboratory Corporation of America) using support from 23andMe.

### Acknowledgments

We are extremely grateful to all the families who took part in this study, the midwives for their help in recruiting them, and the whole ALSPAC team, which includes interviewers, computer and laboratory technicians, clerical workers, research scientists, volunteers, managers, receptionists and nurses.

## Abbreviations

ALSPAC: Avon Longitudinal Study of Parents and Children
ARIES: Accessible Resource for Integrated Epigenomics Studies
BMI: Body mass index
CpG: Cytosine-phosphate-Guanine CRP C-reactive protein
EWAS: epigenome-wide association study
GWAS: genome-wide association study
IVW: inverse-variance weighted
LD: Linkage Disequilibrium
mQTL: methylation quantitative trait loci
MR: Mendelian randomization
PTSD: Post-traumatic stress disorder
QQ: Quantile-quantile
SM: Supplementary Material
SNP: single nucleotide polymorphism
TPM: transcript per million
WM: weighted mean

